# Characterizing patterns of selection pressure on mammalian antiviral immune response

**DOI:** 10.1101/2023.09.26.559440

**Authors:** Mohamed B.F. Hawash

## Abstract

Immune response is known to be under constant pressure evolutionary pressure from different factors including pathogens. Although different selection regimes are expected to act on the magnitude of immune response, there are limited studies that investigated the different patterns of selection pressures on the immune response quantitatively. I employed evolutionary models (Ornstein-Uhlenbeck models) to identify different patterns of selection on the antiviral immune response of fibroblasts derived from 18 mammalian species and one vertebrate stimulated by viral ligand, poly I:C, or Interferon alpha cytokine. I found stabilizing selection to be the dominant form of selection on the immune response. Out of 59 genes that were found to be responding in at least 15 species, 50 genes were found to be under stabilizing selection. Moreover, evolutionary variance was found to differ among these conservatively responding genes implicated in fighting viruses. For instance, ADAR was found to have low evolutionary variance while TRIM14 response showed opposite trend suggesting different evolutionary pressures acting on the magnitude of response. Directional selection was also detected specific infraorders of primates such as apes and old-world monkeys on response of innate immune effectors.

## Introduction

Interspecies comparative studies of immune response have focused mainly on characterizing the conserved immune response across tested species (Barreiro et al., 2010; Gaska et al., 2019; Shaw et al., 2017). While the approach proved to be useful in spotlighting key immune effectors with core functions that are indispensable for host defense, it overlooks other important perspectives such as the relative importance of these conserved genes. For instance, genes that respond with low variability across species are expected to be involved in more core regulatory functions of the immune response unlike genes with higher variability. Moreover, some of the conserved genes may be more responsive in specific clade of the phylogeny revealing directional selection of these genes to perform specific functions in this clade.

In the recent few decades, there has been as expansion of evolutionary models that assess the selection on phenotypic traits while consider the phylogenetic relationship. The main advantage on adopting these models when studying quantitative trait evolution is to incorporate the dependent relationship between species into account (Butler & King, 2004; Cressler et al., 2015). Ornstein-Uhlenbeck (OU) is one the evolutionary models that is designed to distinguish the traits under neutral drift or selection (Butler & King, 2004). The model consists of two component, a stochastic and deterministic parts. The stochastic part pertains to random drift while the deterministic part pertains to selection. Evolutionary models on quantitative trait in the genomic era have focused mainly on gene expression revealing many interesting findings on how gene expression changes contributed to phenotypic differences across species (Catalán et al., 2019; J. Chen et al., 2019). However, there has been a scarcity on application of OU models on other quantitative trait such as immune response. Using same rationale, OU models can disentangle genes whose response is under selection pressure from those that respond in a random drift mode beside genes with directional selection.

Herein, I reanalyzed datasets from interspecies comparative studies of immune response considering the immune response as quantitative trait instead of the original classical approach employed in these studies, conserved genes vs non-conserved. I used an established a statistical framework for testing and applying OU model on quantitative traits developed by Chen and colleagues (J. Chen et al., 2019). My joint analysis is on 19 species from the two studies, one study on primate species (Gaska et al., 2019) and other study on mammalian species (Shaw et al., 2017). I found stabilizing selection to be dominant on the immune response across the 19 species and estimated the evolutionary variance across the conserved genes. Moreover, I found signature of directional selection on old world monkeys and apes that provide new dimensions of the immune response in these clades.

## Materials and methods

### Data collection and genome mapping

I retrieved raw reads from Sequence Read Archive (SRP120495 and ERP023572). The two datasets correspond to two studies that assessed the antiviral response in fibroblasts in primates upon stimulation with PolyI:C, mimicry molecule for viral infection (Gaska et al., 2019) and mammals upon stimulation with IFN alpha cytokine (Shaw et al., 2017) to produce antiviral response. After downloading the data, reads were aligned to respective genomes downloaded from Ensembl. Genome versions used were hg38 for human, Pan_tro_3.0 for chimp, panpan1.1 for bonobo, SaiBol1.0 for Squirrel monkey, gorGor4 for Gorilla, Mmul_10 for rhesus macaque, Panu_3.0 for olive baboon, PPYG2 for orangutan, Mnem_1.0 for pig-tailed macaque, Rnor_6.0 for rat, ARS-UCS1.2 for cow, Oar_v3.1 for sheep, Sscrofa11.1 for pig, EquCab3.0 for horse, CanFam3.1 for dogs, pteVam1 for megabat (large fruit bat) and Myoluc2.0 for microbat (little brown bat). Reads were aligned to genomes using STAR aligner (Dobin et al., 2013) using default parameters. Gene count matrices were generated using featureCounts (Liao et al., 2014). Orthologs for all species were retrieved from Ensembl database v.98. I restricted the orthologous genes used to 1:1 orthologs in all species using human as a reference species. To generate matrices of transcript per million (TPM), I used Kallisto (Bray et al., 2016) to align reads to respective species transcriptomes with default parameters. The R-library tximport (Soneson et al., 2015) were used to summarize TPM values of transcripts to genes.

### Statistical analysis and evolutionary models

I restricted genes for further downstream analysis to those that have more than one count per million (CPM) in at least half of the individuals (>65 samples). All statistical analysis was done on R version 4.2 (https://www.r-project.org/). Count data were normalized using different methods to correct for different library size and sequencing depth using Trimmed Mean of M-values (TMM) method (Robinson & Oshlack, 2010). Scaling factors were calculated by TMM method by estimating the relative fold change of total genes between two libraries under the assumption that majority of genes are not differentially expressed. The ratio of RNA content between the two libraries implements weighted trimmed mean of log ratio where the weights used are the precision (Robinson & Oshlack, 2010). Further normalization was done using voom algorithm (Law et al., 2014) to allow using linear models by limma package. Voom algorithm models mean-variance trend of logCPM for each gene and use it to compute the variance as a weight of logCPM values. “ouch” package version 2.14.1 (Butler & King, 2004) was used to fit ornstein-uhlenbeck (OU) and Brownian motion (BM) models using default parameters. I compared between BM and OU models using likelihood ratio (LR) test for each gene and corrected for multiple testing using Benjamini-Hochenberg (Benjamini & Hochberg, 1995). I fitted OU models with specified optimum in specific subclade (θ_subclade_) and p values were calculated testing OU model with specific clade to OU model with no optimum (θ_subclade_ ≠ θ_ancestral_) and BM model using likelihood ratio test LRT and corrected for multiple testing using FDR (Benjamini & Hochberg, 1995). Akaike Information Criterion (AIC) scores were also used to select for clade specific genes.

## Results

### Conserved antiviral response across species

I analyzed the antiviral response upon stimulation with TLR3 agonist, PloyI:C or IFNA cytokine in the 19 species after 24 hours. I started with 6305 orthologs across all species. After filtering our weakly expressed genes, defined as genes with expression of less than 1 count per million (< 1 CPM) in 65 samples (total replicates from all species are 124), I had 4464 orthologous genes.

I first wanted to characterize the phylogenetic relationship between the tested species. I employed unsupervised clustering method, principal component analysis (PCA) and hierarchical clustering. Samples were clustered primarily according to the phylogenetic relationship between species (figure 1a). The first principal component (PC1) separates apes, monkeys and other mammals while PC2 separates old and new world monkeys from other species (figure 1A). Hierarchical clustering was performed to genes that are responding significantly to stimulants at FDR < 0.15 in at least 9 species (n=428). Samples were also clustered by species that respects phylogenetic relationship then by treatment (Figure 1B,C). Although mouse and rat samples were from the two different studies, they were clustered together suggesting phylogenetic signal to overcome the batch effect (Figure 1A,B). However, large fruit bat was distinctly clustered from other species. This likely be attributed to the constitutive higher expression of antiviral genes as suggested by Shaw and colleagues (2017).

**Figure 1:**
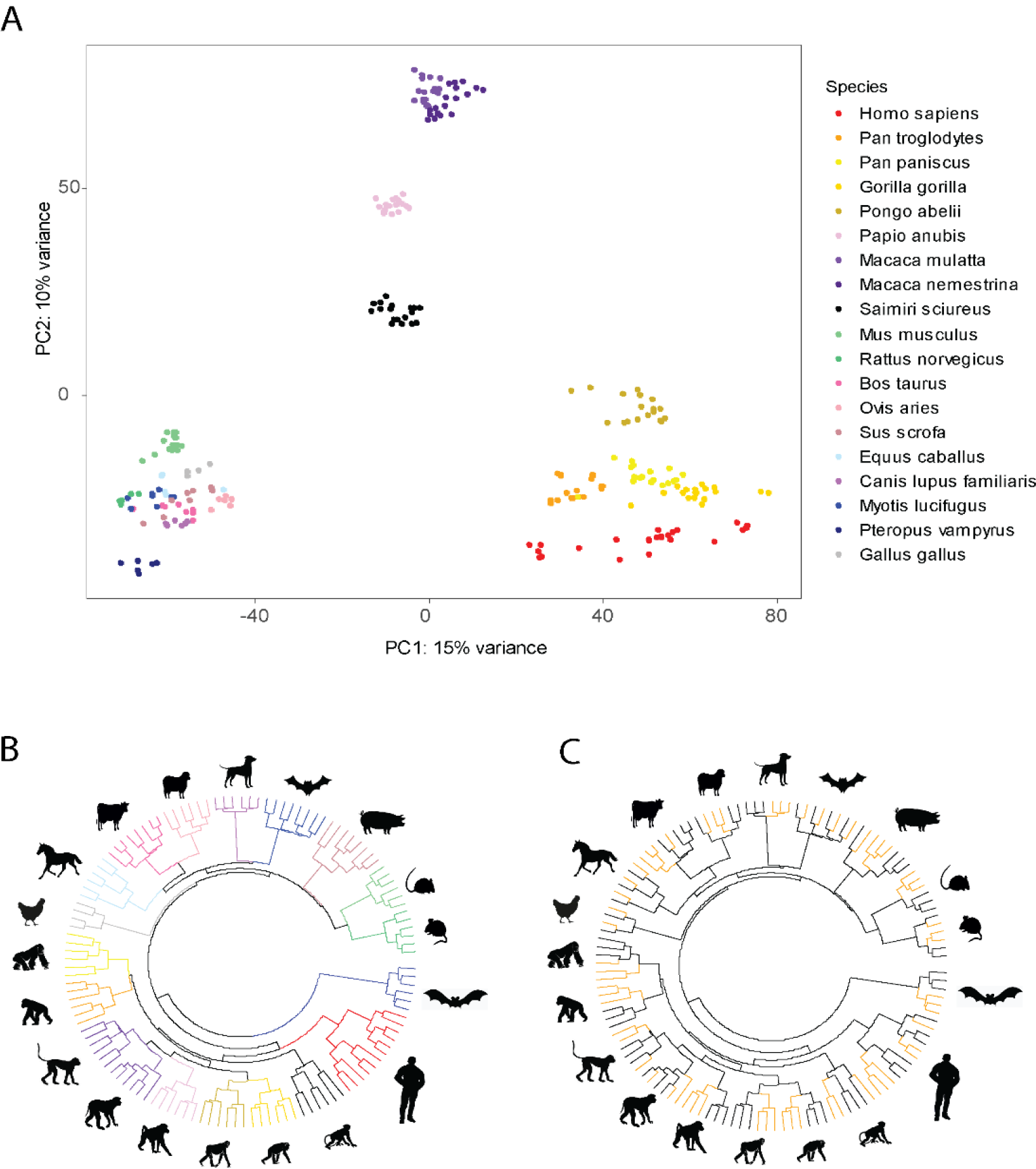
Clustering of Samples respects phylogenetic relationships between species. A. Principal component analysis (PCA) of all samples from all species shows clustering of the samples respects the large know infraorders of mammalian species where PC1 separates apes and mammals and PC2 separates monkeys. B, C. Hierarchical clustering of the genes responding in at least 9 species (n=428) shows clustering of samples follows mostly the know phylogeny of the species; panel C shows the same clustering but highlighting samples without treatment (black line) and samples after viral stimulation (organ line).

Next, I used the linear regression model of ~ species + species:treatment. The number of differentially expressed genes (DEGs) (FDR < 0.15) that were upregulated ranges from 22 to 443 depend on species (Figure S1). Form those, I found that there 15 genes were responding in all species (Table S1). These gene are likely inevitable for the antiviral innate immune response pinpointed by their conservation for nearly 300M years. These genes include antiviral sensor such as TLR3. TLR3 is member of the toll-like receptor mainly involved in recognizing double-stranded RNA viruses (CHEN et al., 2021).

### Patterns of selection on the antiviral innate response

Next, I wanted to characterize which pattern of natural selection involved in shaping the evolution of antiviral response in mammals and primates. I applied OU and BM models and compared between them by LRT. Genes under selection constraint are detected by OU models while ones that respond according to random drift are detected by BM model. The main formula of OU models is dX_t_ = σdB_t_ + α(θ – X_t_)dt where dX_t_ represents the change of trait of interest by time, and dB_t_ is the Brownian motion process. The model calculates 3 major parameters, σ, α and θ. θ represents the optimum to which the trait is pulled at while α is the strength of selection that direct the trait to the optimum θ. σ is the rate of the random change by genetic drift of the trait (J. Chen et al., 2019).

I run the analysis on the two datasets and a joint dataset that comprises both. I found majority of responsive genes (FDR < 0.15, see methods) are under stabilizing selection upon stimulation in joint dataset and primate datasets (Figure 2A, S2). In the joint dataset, I included genes that were significantly upregulated in at least 15 species (FDR < 0.15, n=59). I chose this arbitrary number to ensure that these genes are responding in majority of the species (~ 80% of species number). 50 genes were found to be under stabilizing selection at FDR < 0.05 (Figure 2). There were no significant genes responded in BM vs OU (FDR cutoff 0.15) but the rest of genes that was not significantly under stabilizing selection are depicted in Figure 2B (n=9). These genes close to linear response with respect to genetic distance between species. In the primate dataset, I included significant genes (FDR < 0.15) in at least 7 species (n=451). I found total of 319 genes respond under stabilizing selection in primates and 45 under neutral drift in primates. In the mammalian dataset, I tested 57 genes that are responsive in at least 7 species (FDR < 0.15) and found only 2 genes responded under stabilizing selection. This is perhaps due to the broad evolutionary distance between the mammalian species in this analysis. The signal was detected when the mammalian and primate species included together, the joint dataset, to produce more coherent evolutionary distance between species even though species were treated with different viral stimulants.

**Figure 2.**
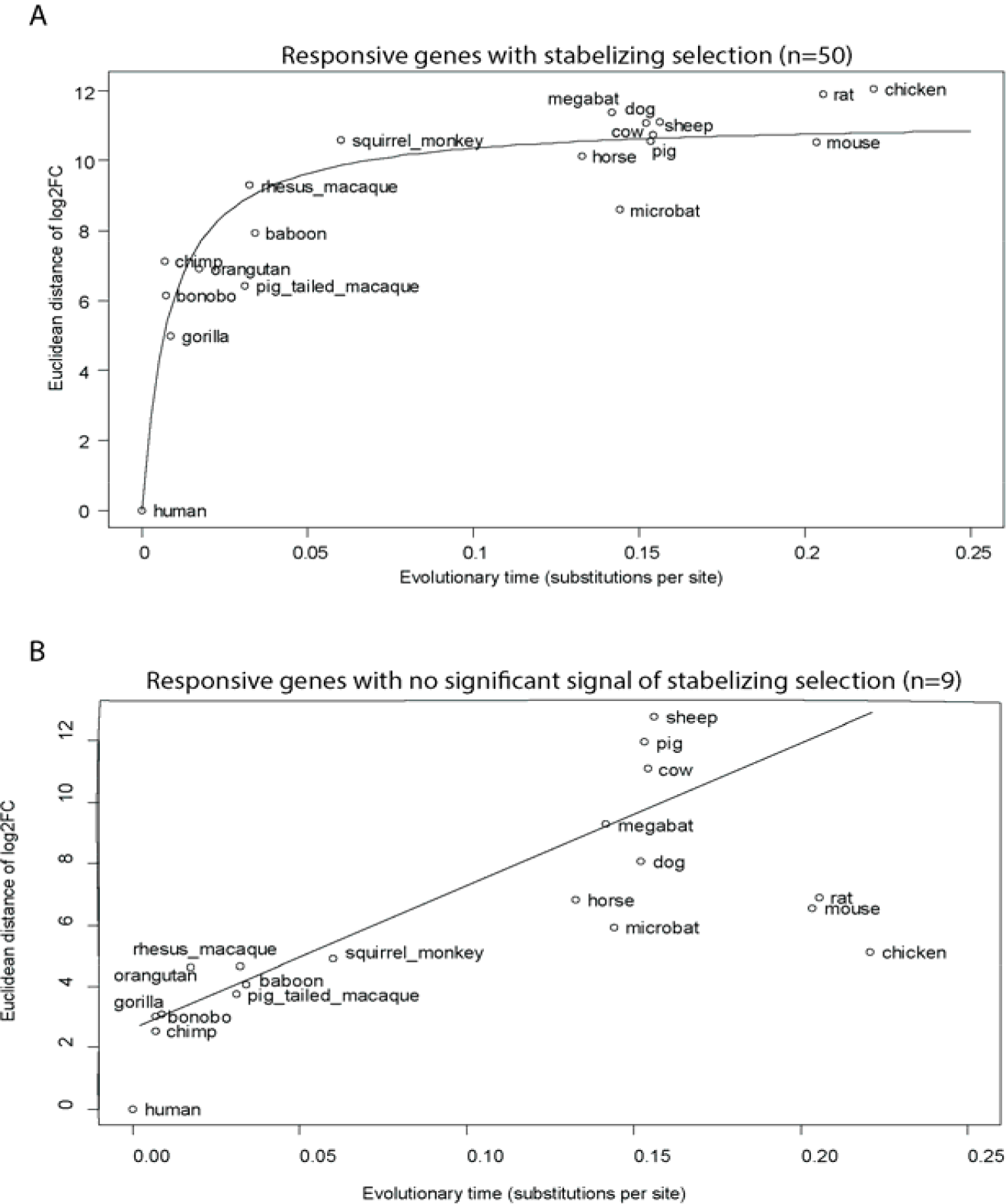
Stabilizing selection shapes the antiviral innate immune genes response in mammals. A. immune responsive genes are under stabilizing selection, i.e., non-linear (n=50) shown by the pairwise differences between human and each species. B. immune genes responding by random drift (n=9).

Next, I wanted to measure the extent of evolutionary constraint of the gene’s responses using “evolutionary variance” parameterized by σ^2^/2α for 59 genes that are significant in >= 15 species. ADAR was among the genes with lowest variance while genes such as TRIM14 had higher variance as shown in Figure 3A,B. Both genes are innate immune regulators that tune the antiviral response through different mechanisms. This discrepancy between the two regulators suggesting there are a secondary layer of selection on the magnitude of response beyond their conserved upregulation. This is remarkable since it suggests there are a single optimum magnitude of response for ADAR for all mammals and the vertebrate species included, chicken, i.e., too much or too little upregulation of ADAR may reduce fitness.

**Figure 3.**
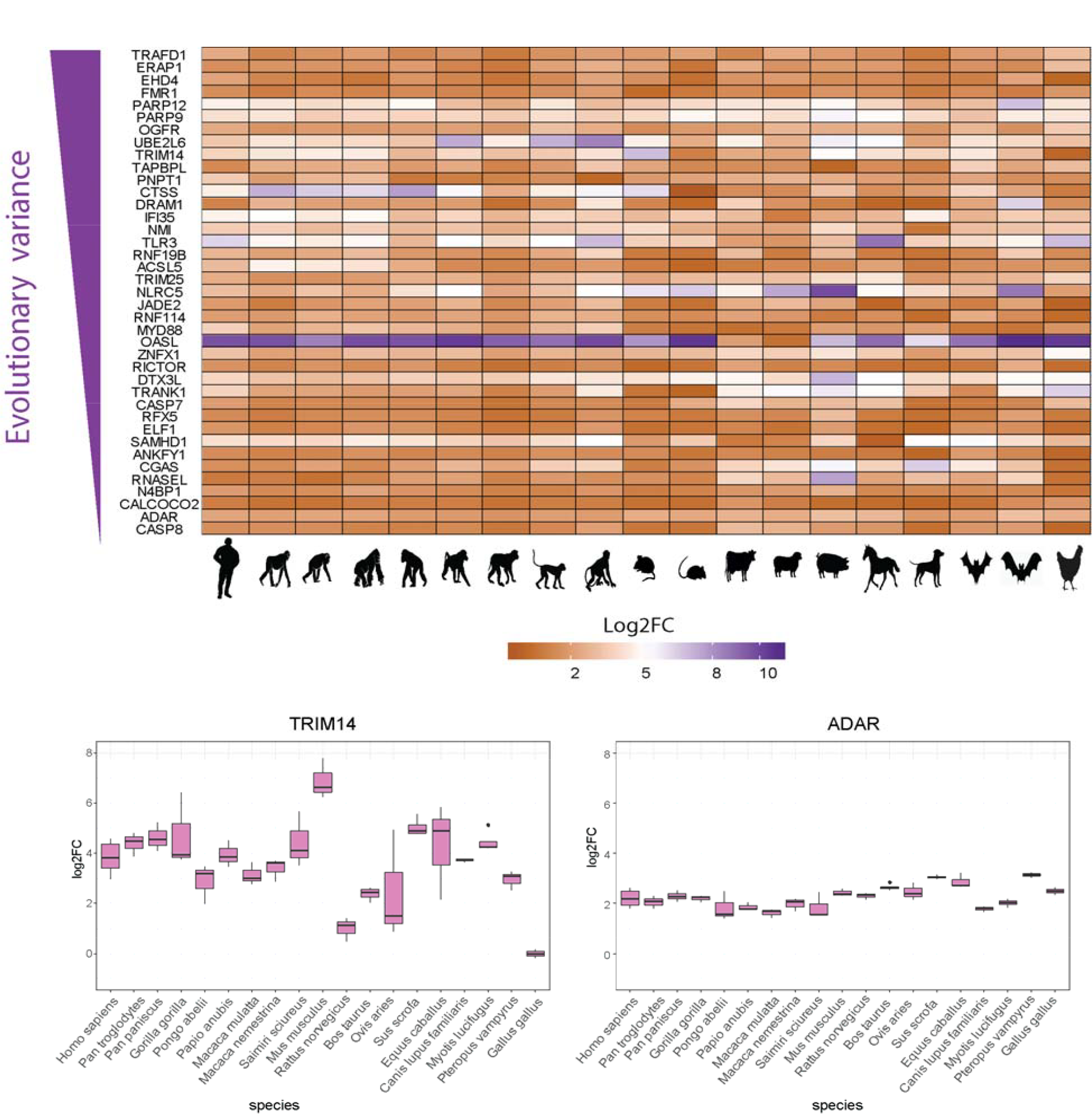
Evolutionary variance of antiviral innate immune genes across mammals. Heatmap of genes responded in at least 15 species ranked according to their estimated evolutionary variance form lowest (bottom) to high variance on top with same examples of ADAR and TRIM14 genes.

Lastly, I wanted to detect the signal of directional selection in specific species clades. I run OU model while specifying optima on 4 clades namely, apes, old world monkeys and rodents and the two bat species. I avoided clades that are completely confounded by study such as all primates or mammals. Apes include human, chimp, bonobo, gorilla and orangutan. Old world monkey clade includes baboo, rhesus macaque and pig-tailed macaque while rodents clade include mouse and rat. I found total of 35 genes were enriched in apes such as PCDH17 and 56 in OWMs such as LXN (Figure 4 A,B). Clade specific genes were significantly different using LRT (FDR < 0.01) and AIC scores differences between the subclade (ape or OWM) and parent primate clade > 10. No significant genes were detected in bats and rodents’ clades. Gene ontology enrichment of these gene. Gene ontology enrichment analysis on the ape-enriched genes identified 32 significant terms (FDR < 0.05) (Table S2). Among the enriched terms “regulation of MAPK cascade” and “respond to other lipopolysaccharide” suggesting that apes are regulating other specific immune pathways when fighting viral pathogens. Similar observation has been reported on apes to be less specific in their immune response (Hawash et al., 2021, 2022). On the other hand, OWM specific genes were enriched for 8 GO terms (FDR < 0.05) (Table S2). Among the enriched Go terms is “response to glucocorticoid”.

**Figure 4.**
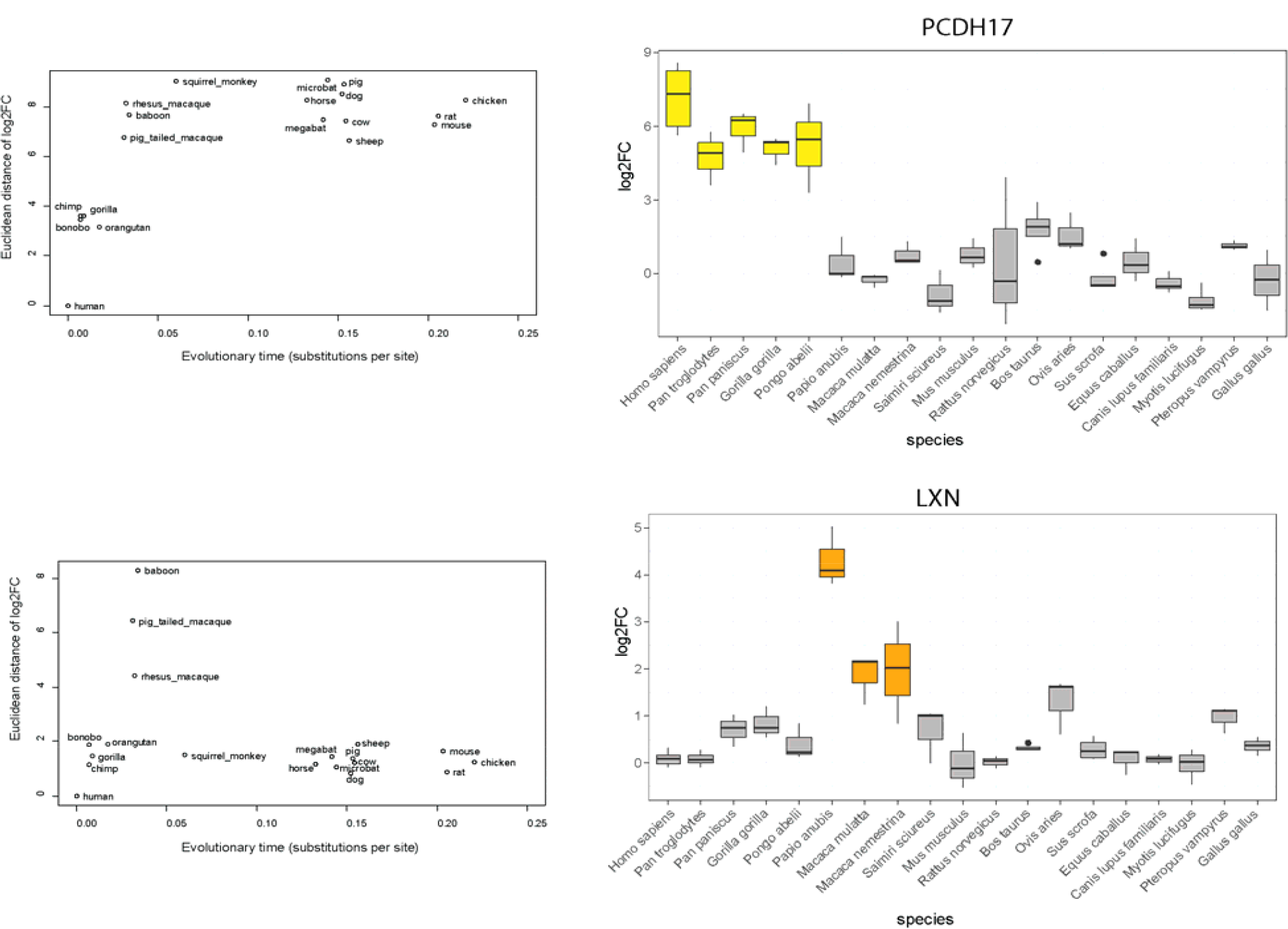
Directional selection in apes and monkeys infraorders. Genes with significant optimum response in apes (top) or old world monkeys (bottom) were characterized. Pairwise distance successfully identified the clade specific genes (left side) and examples are shown on the right. Gene lists and significantly enriched GO terms for these genes are given in Table S2.

Glucocorticoid is a group of hormones that alleviate the inflammation and diseases including cancer (Timmermans et al., 2019). Hence, the OWMs response may therefore aim to mainly alleviate the collateral damage resulted from the inflammation and the harsh innate immune response.

## Discussion

In this study, I aimed to characterize the pattern of selection on the immune response as a quantitative trait. The advantage on having an evolutionary statistical framework to study the immune response is to determine the extend of constrain on the response. The classic approach in comparative analysis across species immune response is to determine the conserved genes. While this approach proved useful in characterizing genes indispensable for species immune response, but it misses the information on how much constrain is exerted on specific genes. For instance, I found low evolutionary variance on ADAR unlike TRIM14 suggesting higher constrain on ADAR response. ADAR, also know as ADAR1, is well know regulator of antiviral immunity. ADAR1 is a highly conserved proteins in animals and it is mainly responsible for modifying the adenosine sites in dsRNA to inosine (A-I) deamination (Quin et al., 2021). This process is to distinguish the self RNA molecules form foreign molecules to avoid triggering innate immunity by self-molecules and it mediated by specific form of ADAR1 known as ADAR1p110. Another form of ADAR1, ADAR1p150 is involved in antiviral response by activation of innate immune receptors know as RIG-I– like receptors (RLR). Activation of ADAR1p150 can edit any RNA including RNA making it similar to self RNA molecules, thus, enable evasion of viruses form antiviral restriction (Quin et al., 2021). Hence, it makes sense that the expression is under strict upregulation in all species tests since although it leads to activation of antiviral pathways, but it also may enable immune evasion of viruses necessitating strict control on its activation. On the other hand, TRIM14 has been found to be implicated in type I interferon (IFN) pathway that regulate interferon stimulating genes (ISGs) upon viral or bacterial stimulation (M. Chen et al., 2016; Hoffpauir et al., 2020). TRIM14 was reported to activate the type I IFN response upon virus infection as evidenced by susceptibility of TRIM14 knock out mice to herpes simplex virus infection (M. Chen et al., 2016). In contrary, TRIM14 was found to found to downregulate type I IFN upon bacterial stimulation with *Mycobacterium tuberculosis* in mouse macrophages (Hoffpauir et al., 2020). Moreover, TRIM14 knock out macrophages was found also to upregulate more antimicrobial peptides suggesting overall negative effect of TRIM14 on immune response. TRIM14 is therefore likely plays different roles in the immune response that is context dependent and thus its response fluctuates across species. ADAR1 is essential for survival of mouse as indicated by knockout mice that die at embryonic day 11.5-14 (Quin et al., 2021) while no such know phenotype for TRIM14.

Although both ADAR and TRIM14 are regulators of antiviral immune response that are induced by majority of mammalian species, their level of induction provides more information on their relative functional importance. Genes such as ADAR with low variability across species are likely to be involved in core process of the innate immune response that mandates strict and fine-tune upregulation. On the other hand, genes such as TRIM14 with high variability across species are likely to be involved in different mechanism of immune response.

In summary, modelling of innate immune response as a quantitative trait will provide further insights on the functional importance of effectors beyond the simple binary approach widely adopted. I urge to exploit the interspecies comparative datasets of immune response with similar quantitative approach to reveal other important features of the immune response across species.

## Statements and Declarations

### Funding

The author declares that no funds, grants, or other support were received for the submitted work

### Competing Interests

The author declares no competing interests.

### Ethical approval

This is meta-analysis paper and the ethical approval was provided in the original studies.

### Availability of data and materials

The data used in this study is available in the Sequence Read Archive under accession numbers SRP120495 and ERP023572

## Notes

### Competing Interest Statement

The authors have declared no competing interest.

